# ATP-induced crosslinking of a biomolecular condensate

**DOI:** 10.1101/2023.04.18.535486

**Authors:** Sebastian Coupe, Nikta Fakhri

## Abstract

DEAD-box helicases are important regulators of biomolecular condensates. However, the mechanisms through which these enzymes affect the dynamics of biomolecular condensates have not been systematically explored. Here, we demonstrate the mechanism by which mutation of a DEAD-box helicase’s catalytic core alters ribonucleoprotein condensate dynamics in the presence of ATP. Through altering RNA length within the system, we are able to attribute the altered biomolecular dynamics and material properties to physical crosslinking of RNA facilitated by the mutant helicase. These results suggest the mutant condensates approach a gel transition when RNA length is increased to lengths comparable to eukaryotic mRNA. Lastly, we show that this crosslinking effect is tunable with ATP concentration, uncovering a system whose RNA mobility and material properties vary with enzyme activity. More generally, these findings point to a fundamental mechanism for modulating condensate dynamics and emergent material properties through nonequilibrium, molecular-scale interactions.

**Significance:** Biomolecular condensates are membraneless organelles which organize cellular biochemistry. These structures have a diversity of material properties and dynamics which are crucial to their function. How condensate properties are determined by biomolecular interactions and enzyme activity remain open questions. DEAD-box helicases have been identified as central regulators of many protein-RNA condensates, though their specific mechanistic roles are ill-defined. In this work, we demonstrate that a DEAD-box helicase mutation crosslinks condensate RNA in an ATP-dependent fashion via protein-RNA clamping. Protein and RNA diffusion can be tuned with ATP concentration, corresponding to an order of magnitude change in condensate viscosity. These findings expand our understanding of control points for cellular biomolecular condensates that have implications for medicine and bioengineering.

## 1 Introduction

Biomolecular condensation is a form of cellular organization which has been implicated in a wide variety of cellular processes including embryonic development, ribonucleoprotein complex assembly, stress response, and misfolded protein storage [1]. The biochemical roles of biomolecular condensates are thought to include locally concentrating reactions, sequestration of biomolecules, and cytosolic concentration buffering [1][2]. These structures form when biopolymers, often a combination of RNA and protein, undergo a phase transition into a more concentrated liquid-like or gel-like state [3]. Specific protein-protein, protein-RNA, and RNA-RNA interactions within these memebraneless organelles produce micro-structures which affect dynamics within biomolecular condensates [4, 5]. Biomolecular condensate micro-structure and dynamics are expected to define their reaction kinetics, molecular size filtering, and composition [2]. Increased understanding of the connection between the molecular interactions of condensate components and condensate structure and dynamics is fundamental to understanding their biochemical role within the cell and also key to their application in bioengineering.

One class of proteins, the DEAD-box helicase, has gained recent appreciation for the role it may play in regulating biomolecular condensate structure and dynamics [6–8]. DEAD-box helicases remodel RNA-RNA and RNA-protein interactions in an ATP-dependent manner [9, 10]. DEAD-box helicases have been identified as important for a variety of cellular processes such as mRNA nuclear export, RNA splicing, mRNA cap recognition, and ribosome assembly [10]. They are also believed to regulate condensate formation through their enzyme activity and they may additionally be capable of regulating the material state of condensates [6]. Mutating DEAD-box helicases within cells often results in aberrant host condensate localization, morphology, and dynamics [11–13]. Furthermore, mutations associated with human DEAD-box helicase DDX3 driven diseases such as medullablastoma and DDX3 syndrome also promote abberant stress granule formation in human cells [14, 15]. An open question remains whether this class of proteins impacts condensate dynamics via their interactions with single-stranded RNA, unwinding duplexed RNA, or a combination of these two activities. DEAD-box helicase ssRNA clamping has been proposed to induce gel-like condensates based on in vivo experiments, though we still lack direct evidence for this phenomenon beyond disrupted biomolecular dynamics [6, 7, 11, 16].

We address the role of ATP-dependent ssRNA binding using a model condensate forming DEAD-box helicase, LAF-1. The LAF-1 protein is a *C. elegans* DEAD-box helicase orthologous to human DDX3, *S. cerevisiae* Ded1p, and *D. melanogaster* Belle, and localizes to the P granule condensate in *C. elegans* germ cells and embryos [17, 18]. LAF-1 can phase separate in vitro to form condensates with liquid-like properties which incorporate RNA and whose properties depend on RNA length [19–21]. However the influence of LAF-1 enzymatic activity and nucleotide-binding on the material properties of the condensates it forms remains unstudied.

To understand the role of LAF-1’s ATP hydrolysis in determining condensate dynamics, we characterize how a LAF-1 mutant alters condensate biomolecular dynamics and material properties in the presence of ATP and single-stranded RNA. Our results suggest the effect on condensate dynamics is the result of the time it spends in the midst of its ATPase cycle as opposed to the lower overall ATP turnover. We then gain mechanistic insight by comparing changes in protein versus RNA dynamics within the condensed phase. Through varying RNA length, we observe a dramatic increase in effective viscosity for the mutant helicase relative to the wildtype enzyme which correlates with a slowdown in RNA mobility and changes in protein diffusion mode, strongly implying a crosslinking mechanism. Lastly, we show the effects on material properties and biomolecular dynamics are titratable with ATP concentration, with a transition point consistent with the helicase’s ATP binding thermodynamics. These results both demonstrate a mode by which a commonly studied DEAD-box mutation affects condensate properties and shed light on a general mechanism for generating micro-scale structure within biomolecular condensates via interaction lifetimes, relevant for both bioengineering and human health and disease.

## 2 Results and Discussion

### 2.1 A Mutant Trapped in an ATP-Bound Intermediate Alters Condensate Dynamics and Material Properties

While LAF-1 has been shown to phase separate in the presence or absence of RNA, the behavior of LAF-1 condensates in the presence of nucleotide has not been previously described [19, 21]. To assess how nucleotides and ATP hydrolysis affects the microenvironment of LAF-1 condensates containing RNA, we purified LAF-1^*WT*^ and two LAF-1 mutants whose mutations within the catalytic ATPase domain have been studied for their effects on ATP binding and hydrolysis in other DEAD-box helicases. LAF-1 (E398A), LAF-1^*DAAD*^, was expected to have decreased ATPase activity through an inability to efficiently enter into its ATPase cycle [16, 22, 23]. LAF-1 (E398Q), LAF-1^*DQAD*^, was expected to have severely impeded ATP hydrolysis activity due to an inability to release the products of ATP hydrolysis, resulting in the formation of long-lived LAF-1^*DQAD*^-RNA complexes in the presence of ATP [11, 24]. We confirmed decreased ATPase activity in these mutants via a kinetic assay measuring phosphate release in the presence of ssRNA (Figure S1). All three variants are capable of forming condensates in the presence of the single-stranded RNA (ssRNA) poly-uridylic acid (PolyU) and ATP or ADP.

To assess whether the LAF-1 catalytic substitutions affected condensate dynamics, we utilized fluorescence recovery after photobleaching (FRAP). Condensates with fluorescently labelled PolyU RNA and protein were prepared in the presence of ADP or ATP (Figure 1c). Small regions composing *≤* 20% of the condensate volume were photobleached, and the recovery dynamics measured (Figure 1d). Recovery curves were normalized relative to initial droplet intensity, bleach spot intensity, and droplet intensity far from the bleach spot (Supplementary Information) [25]. All three LAF-1 variants exhibit similar protein FRAP behavior in the presence of ADP (Figure S2a). When ATP is present instead, the LAF-1^*WT*^ and LAF-1^*DAAD*^ variants behave similarly to one another and to their counterpart systems with ADP (Figure 1e). However, the LAF-1^*DQAD*^ system exhibits a drastic change in protein dynamics in the presence of PolyU and ATP (Figure 1e). At short timescales, there is fluorescence recovery with a similar timescale to the other variants, but to a much lesser extent. There is also a slower recovery timescale that allows the LAF-1^*DQAD*^ protein to recover to roughly same final extent as the other variants. Protein recovery dynamics for all variants in the presence of ADP were fit to a single exponential, as were the dynamics of LAF-1^*WT*^ and LAF-1^*DAAD*^ in the presence of ATP (Figure 1f). For the LAF-1^*DQAD*^ variant with ATP, we fit the protein recovery dynamics to a double exponential to capture both recovery timescales (Figure 1f). The short timescale recovery dynamics of the LAF-1^*DQAD*^ system with ATP matches the timescale of the other variants with either nucleotide. The second, longer timescale is over an order of magnitude greater than this fast timescale behavior. The emergence of a second timescale in the LAF-1^*DQAD*^ FRAP raises the possibility that this slower state corresponds to its clamped state on RNA.

**Figure 1.**
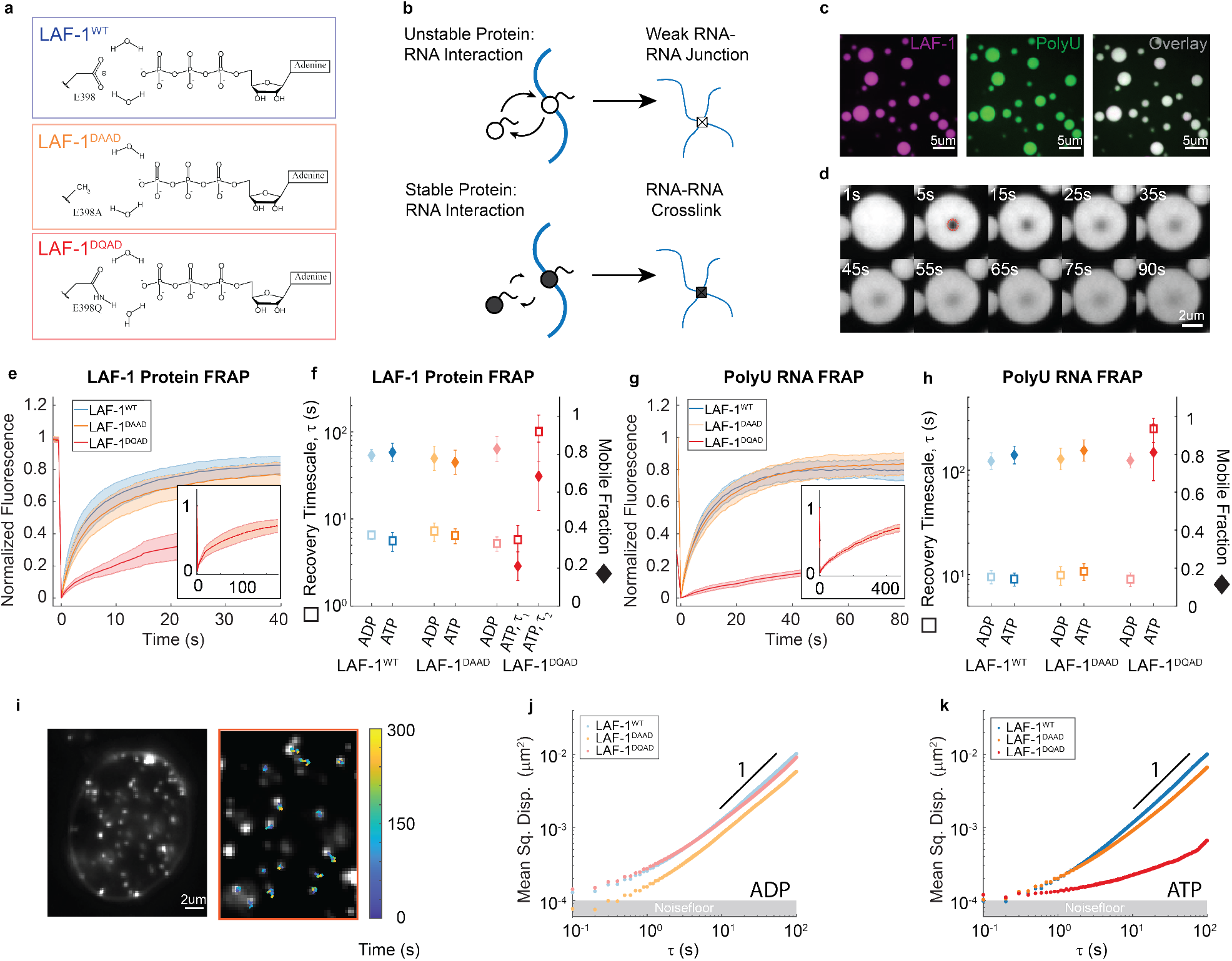
Trapping an intermediate along the ATP-hydrolysis pathway induces slowed dynamics and altered material properties of a DEAD-box helicase-RNA condensate. a) Mutation scheme for the LAF-1 DEAD-box helicase using well characterized DEAD-box mutations. b) Transient LAF-1-RNA association does not enable strong RNA-RNA interactions but long-lived protein-RNA clamping results in RNA-RNA crosslinking. c) LAF-1 condensates co-condense with PolyU RNA in the presence of ATP. d) Representative fluorescence recovery after photobleaching (FRAP) assay to study macromolecular diffusion in biomolecular condensates. e) FRAP curves and f) extracted fit parameters for LAF-1 protein in LAF-1-PolyU condensates indicate that clamp-like mutant LAF-1^*DQAD*^ but not slowed ATPase mutant LAF-1^*DAAD*^ experiences altered dynamics in the presence of ATP. Two timescales are apparent in LAF-1^*DQAD*^ recovery, with the faster timescale matching that of LAF-1^*WT*^. g) FRAP curves and h) extracted fit parameters for PolyU RNA in LAF-1-PolyU condensates indicate over an order of magnitude decrease in RNA diffusion in clamp-like LAF-1^*DQAD*^ condensates in the presence of ATP. i) Tracking of fluorescent tracer particles in biomolecular condensates to measure rheological properties j) and k) Mean squared displacement plots for beads embedded in LAF-1-PolyU condensates indicate altered material properties for clamp-like mutant LAF-1^*DQAD*^ in the presence of ATP, with diffusive, fluid-like dynamics for all other conditions.

If the second timescale in the LAF-1^*DQAD*^ protein FRAP data corresponds to the clamped state, it should track with the diffusion of RNA within the system. We next examined the FRAP response of labelled PolyU RNA within the different LAF-1 systems in the presence of ATP or ADP (Figure 1g and Figure S2b). We fit the curves to single exponentials and extracted recovery timescale and mobile fraction fit parameters (Figure 1j). LAF-1^*WT*^ and LAF-1^*DAAD*^ have similar RNA FRAP responses in the presence of ATP and ADP, and are similar to one another. LAF-1^*DQAD*^ has a similar RNA FRAP response to the other variants in the presence of ADP. However, when ATP is present, the recovery timescale increases by over an order of magnitude (ADP: 9.0s, ATP: 250s) while the RNA mobile fraction remained roughly the same (ADP: 0.77, ATP: 0.80). This single timescale visible in the RNA recovery dynamics is roughly comparable to the second, longer timescale visible in the LAF-1^*DQAD*^ protein recovery dynamics suggesting the slow protein timescale corresponds to a LAF-1^*DQAD*^ species clamped to RNA.

Though both LAF-1^*DAAD*^ and LAF-1^*DQAD*^ have impeded ATPase activity, only LAF-1^*DQAD*^ experiences any change in biomolecular dynamics, and only when in the presence of ATP. This indicates that the changes in biomolecular dynamics are likely due to the trapped conformation of LAF-1^*DQAD*^ bound to RNA and ATP hydrolysis products, as opposed to simply decreased ATPase activity [11]. Furthermore, the appearance of a second timescale within the LAF-1^*DQAD*^ diffusion dynamics in the presence of ATP, which matches the RNA diffusion timescale of the system, also suggests that the LAF-1^*DQAD*^ protein might be travelling with the RNA in this slower diffusion mode. The significant slowdown of RNA mobility within the LAF-1^*DQAD*^ condensates also poses the possibility that the LAF-1^*DQAD*^ proteins trapped on the RNA may be acting as RNA crosslinkers within the system.

If crosslinking is occurring, there should be corresponding changes in the material properties of LAF-1^*DQAD*^ condensates. Point tracking microrheology has been used to study biomolecular condensate rheology previously, including for the LAF-1 system [19–21, 26]. We embedded PEG-passivated fluorescent tracer particles with a 100nm diameter in LAF-1 condensates during condensate formation, and measured bead positions over time (Figure 1i). From these trajectories, we calculated the mean squared displacement (MSD) of the beads and took a time-ensemble average over beads within individual droplets (Supplementary Information). In this way, we measured MSDs for beads in condensates of each of the three LAF-1 variants with PolyU RNA and ADP or ATP (Figure 1l and 1m). Beads in condensates of all three variants with PolyU RNA and ADP behave quite similarly (Figure 1j). The curves are diffusive in the long time limit, consistent with fluid-like behavior, and the subdiffusive behavior at short times can be accounted for through a constant noisefloor contribution (Supplementary Information). When ATP is present, though beads in the LAF-1^*WT*^ and LAF-1^*DAAD*^ behave similarly and diffusively, bead dynamics within the LAF-1^*DQAD*^ system are effectively arrested over the course of their trajectories (Figure 1k). The slowly hydrolyzable ATP analog ATP*γ*S does not elicit any difference in material properties for LAF-1^*WT*^ when compared to ADP or ATP, providing an additional indication that ATPase activity itself is not responsible for the changes in condensate material properties seen with LAF-1^*DQAD*^ (Figure S3). Thus the effects seen at the macromolecular scale via FRAP are recapitulated in a substantial change in the material properties of LAF-1^*DQAD*^ condensates.

### 2.2 RNA Length Dependent Material Properties Reveals Hydrogel-like Properties of LAF-1^*DQAD*^ Condensates

We next sought to gain better insight into the material effects of the LAF-1^*DQAD*^ mutation. Because varying concentration of biomolecules in condensates is not straightforward, due to the tendency for liquid-liquid phase separated systems to buffer concentrations, many of the traditional concentration-based experiments to probe polymer-based materials were not possible. Instead, we systematically varied RNA length in order to assess scaling relationships between RNA length and condensate material state. The RNA size distribution for the initial polydisperse PolyU sample was measured via capillary electrophoresis (Figure 2a). We fragmented the PolyU sample using magnesium and heat to generate samples with progressively shorter mean fragment sizes (Figure 2a). Point tracking microrheology experiments were conducted using the resulting RNA samples, keeping the weight fraction of RNA constant for all conditions. We then compared MSDs for beads inside LAF-1^*WT*^ and LAF-1^*DQAD*^ condensates in the presence of 5mM ATP (Figure 2b and Figure 2c). Beads within the wildtype condensate do not experience a significant change in their MSDs for the shorter length distributions, but at the longest mean RNA length the MSD is lower at all time-lags. The MSDs for all lengths are well described by a power law scaling exponent of 1. For all RNA lengths in the LAF-1^*DQAD*^ mutant system with ATP, the bead MSDs are slower than their counterparts in the LAF-1^*WT*^ condensates and also much more sensitive to RNA length. While the beads exhibit diffusive motion in LAF-1^*DQAD*^ condensates with the four shortest RNA length distributions, the longest RNA length sample results in severely slowed bead dynamics where positional fluctuations are indistinguishable from noise. The slower bead dynamics and higher length sensitivity of bead MSDs in the LAF-1^*DQAD*^ system suggested that the underlying material state of the system was changing in response to crosslinking by LAF-1^*DQAD*^.

**Figure 2.**
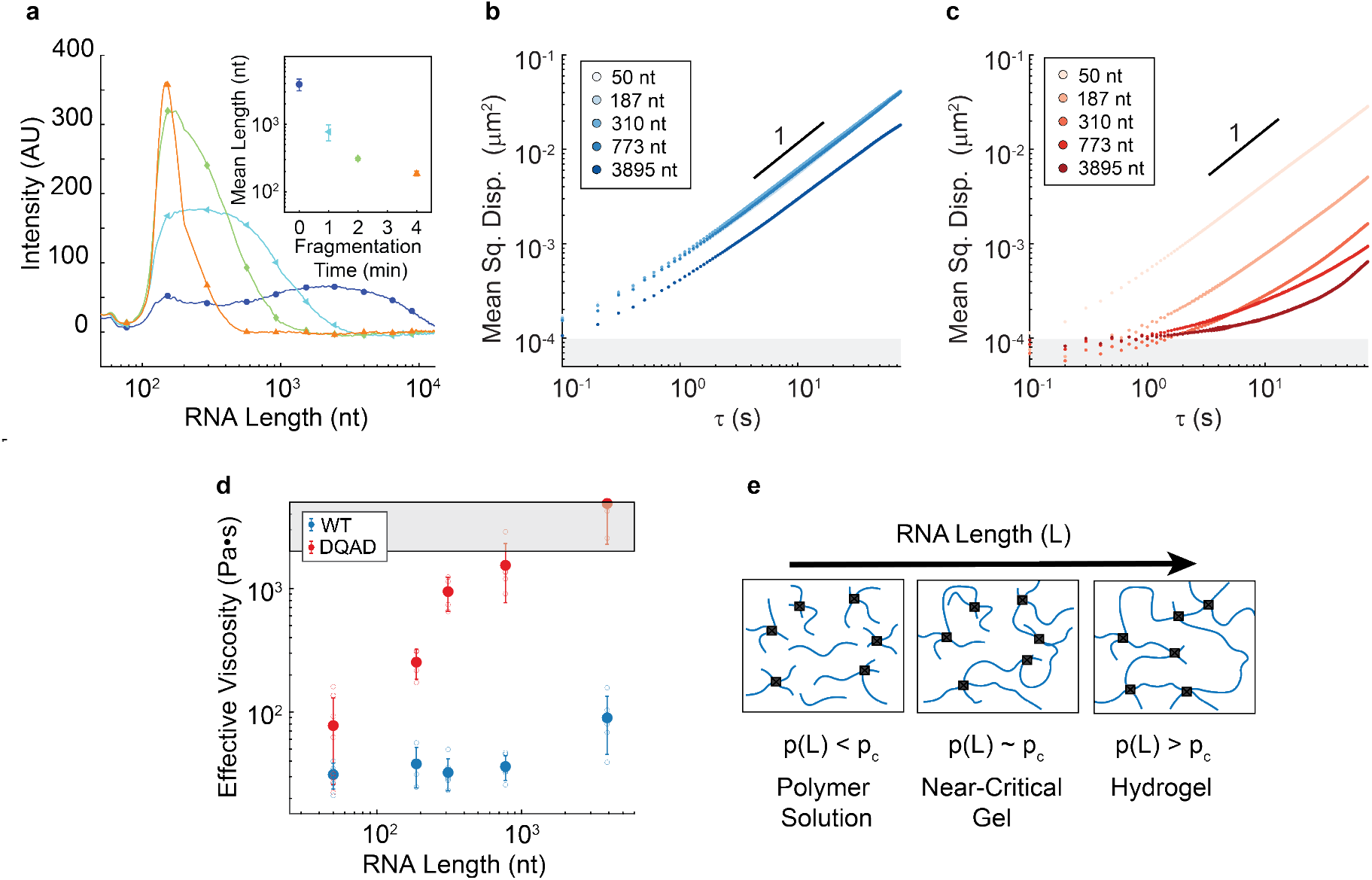
RNA Length-Viscosity Dependence Indicates RNA Crosslinking within LAF-1^*DQAD*^ Condensates. a) Capillary electrophoresis results of heat and magnesium fragmented PolyU RNA for various times. b) and c) MSDs of fluorescent tracer particles in LAF-1^*WT*^ and LAF-1^*DQAD*^ condensates have different responses to RNA length in the presence of ATP. d) Effective viscosity for condensates containing LAF-1^*WT*^ or LAF-1^*DQAD*^ (blue and red, respectively), PolyU RNA of the indicated length, and 5mM ATP, as extracted using the Stokes-Einstein relation. A sharp increase in effective viscosity for LAF-1^*DQAD*^ condensates is observed to begin at a RNA length of roughly 250 nts, consistent with near-critical-gel-like behavior [27, 28]. e) RNA length dependent crosslinking, wherein longer RNAs result in a exponentially larger complexes. The system should eventually tend toward a percolated gel-like RNA network as a critical RNA chain length is reached [29, 30].

To substantiate that the material state of the LAF-1^*DQAD*^ system differed from LAF-1^*WT*^ in the presence of ATP, we assessed the relationship between RNA length and viscosity. For those MSDs that exhibited diffusive scaling with time-lag, we extracted effective viscosities using the Generalized Stokes Einstein Relation (Supplementary Information). Though the systems with ATP are not in equilibrium, Van Hove correlation plots and velocity autocorrelation plots do not suggest active transport of the passive particles is occurring and thus we believe extraction of an effective viscosity is appropriate (Figure S4). A sharp increase in effective viscosity occurs for LAF-1^*DQAD*^ for condensates containing RNA lengths longer than 250 nucleotides (Figure 2d). Both variants produce condensates with similar RNA-length-dependent effective viscosities in the absence of ATP (Figure S5). We observe no substantial changes in partitioning of biological macromolecules as a function of RNA length or ATP concentration (Figure S6). The sharp increase in effective viscosity as a function of RNA length (Figure 2d) is consistent with a near critical gel. Longer RNA lengths should support the formation of exponentially larger complexes in the presence of crosslinking. In such systems, as increasing cluster size begins to approach the percolation threshold for the system, a corresponding dramatic increase in viscosity is observed [27, 28]. We cannot rule out the possibility of larger crosslinked protein-RNA complexes becoming entangled as the source of the jump in effective viscosity. However, this is not necessarily incompatible with our model of RNA-RNA crosslinking facilitated by LAF-1^*DQAD*^ and would similarly result in hydrogel-like material properties [31].

### 2.3 RNA Length Dependent Biomolecular Dynamics Indicates Formation of Supramolecular Complexes

We next wanted to examine the molecular basis for crosslinking by examining biomolecular diffusion within the condensates as a function of RNA length. We measured FRAP curves for labelled LAF-1 protein (Figure 3 a and b) and end-labelled PolyU RNA (Figure 3 d and e) in either LAF-1^*WT*^ or LAF-1^*DQAD*^ condensates using the different RNA fragment distributions. For LAF-1^*WT*^ protein FRAP, the recovery curves are well fit by a single exponential and the timescale of recovery is shown in Figure 3d. For LAF-1^*DQAD*^ protein FRAP, two timescales are apparent in the recovery dynamics and we fit the curves to a double exponential. The shorter fit timescale matches well with that of the LAF-1^*WT*^ protein and was insensitive to RNA length (Figure 3d). The second timescale is larger by at least an order of magnitude from the first and increases with increasing RNA length. This is consistent with a model whereby a portion of the population of the LAF-1^*DQAD*^ molecules are diffusing freely, resulting in a timescale similar to the LAF-1^*WT*^ molecules. The second population of LAF-1^*DQAD*^ forms a long lived complex with RNA such that this second timescale is a combination of the RNA diffusion timescale and the release timescale off of the RNA. This second timescale is thus expected to increase with RNA length, consistent with our data (Figure 3g).

**Figure 3.**
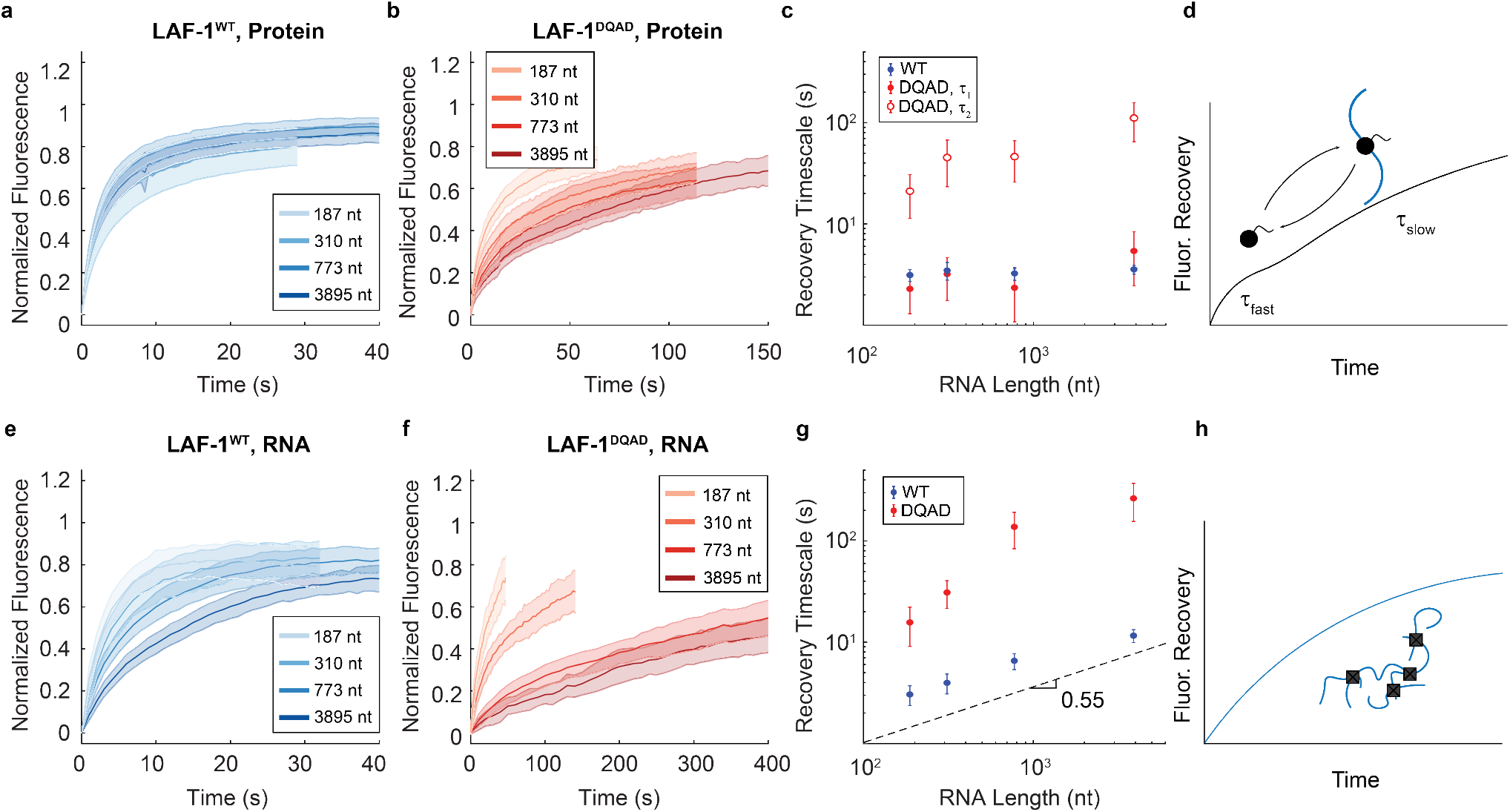
Increased Effective Cluster Size for RNA in LAF-1^*DQAD*^ Condensates. a) and b) LAF-1 protein FRAP curves for LAF-1^*WT*^ and LAF-1^*DQAD*^ condensates, respectively, with varying PolyU lengths and 5 mM ATP. c) FRAP recovery timescales for LAF-1 protein FRAP from (a) and (b). Good agreement is seen between the LAF-1^*WT*^ recovery timescale and the first timescale of the LAF-1^*DQAD*^ fit, with no significant dependence on RNA length. LAF-1^*DQAD*^’s second recovery timescale scales with RNA length. d) LAF-1^*DQAD*^ FRAP curves are best fit by a double exponential. The free, diffusing protein species has a fast recovery timescale while the protein bound to the supramolecular protein-RNA complex will have a slower timescale that is a combination of the network recovery timescale and protein release timescale. e) and f) FRAP curves for labelled PolyU RNA in LAF-1^*WT*^ and LAF-1^*DQAD*^ condensates, respectively, with varying PolyU lengths and 5 mM ATP. g) RNA FRAP recovery timescales fit to single exponentials. A stronger RNA length dependence on RNA recovery timescale is seen for the LAF-1^*DQAD*^ mutant, consistent with the formation of supramolecular ribonucleoprotein complexes. The dotted line represents the expected scaling of recovery timescale with length based on predicted polymer size [32]. h) The RNA will have a single timescale which is either the average recovery of the supramolecular protein-RNA structures or the network’s recovery timescale.

We expected that the RNA length dependence of the slower timescale in LAF-1^*DQAD*^’s diffusion should then correlate with RNA diffusion timescales. We measured the RNA FRAP timescales for the different fragment lengths in LAF-1^*WT*^ and LAF-1^*DQAD*^ condensates in the presence of ATP (Figure 3 d and e). The curves were fit to single exponentials, yielding recovery timescales which increase with RNA length for both variants (Figure 3 f). However, the scaling relationship between length and RNA recovery timescale is much higher for the LAF-1^*DQAD*^ versus LAF-1^*WT*^. The scaling relationship for the wildtype is roughly that which would be expected for a dilute solution of RNA within the protein condensate [32]. The larger scaling relationship for the LAF-1^*DQAD*^ mutant is consistent with the formation of supramolecular complexes due to protein facilitated crosslinking, such that the effective biomolecular size is larger than would be expected given the input molecules (Figure 3h). Importantly, the RNA FRAP timescales roughly match the slower timescale in the LAF-1^*DQAD*^ protein FRAP, supporting the idea that the slower timescale in the protein FRAP corresponds to the RNA-clamped state of the protein. The plateau in the RNA FRAP timescale at very long RNA lengths could indicate that crosslinking of the entire system is accomplished above average RNA lengths of roughly 500 nucleotides and that the timescales above this point measure rearrangement timescales of the resulting gel.

### 2.4 Titratable Crosslinking with ATP Concentration in LAF-1^*DQAD*^ Condensates

Because the properties of the LAF-1^*DQAD*^ were strongly dependent on the presence of ATP, we next determined whether titrating ATP concentration within the system produced tunable system properties. We first tested the ATP concentration-dependent FRAP properties of LAF-1 protein and full-length PolyU RNA for the LAF-1^*DQAD*^ variant. Protein FRAP curves for LAF-1^*DQAD*^ and full length PolyU RNA as a function of ATP concentration are shown in Figure 4a. For MgATP concentrations above 0 mM the curves were fit to double exponentials, with the extracted recovery timescales and mobile fractions as a function of MgATP concentration shown in 4c and d. For the shorter timescale, representing the freely diffusing protein within the condensate, the recovery timescale remains approximately constant over the range of MgATP concentrations tested. This shorter timescale also matches well with the single-exponential fit timescale of the 0mM MgATP condition. The second, longer timescale in the FRAP dynamics increases monotonically with ATP concentration. As ATP is titrated into the system, more LAF-1^*DQAD*^ becomes trapped onto RNA, resulting in larger RNA clusters and therefore a slower second FRAP recovery timescale.

**Figure 4.**
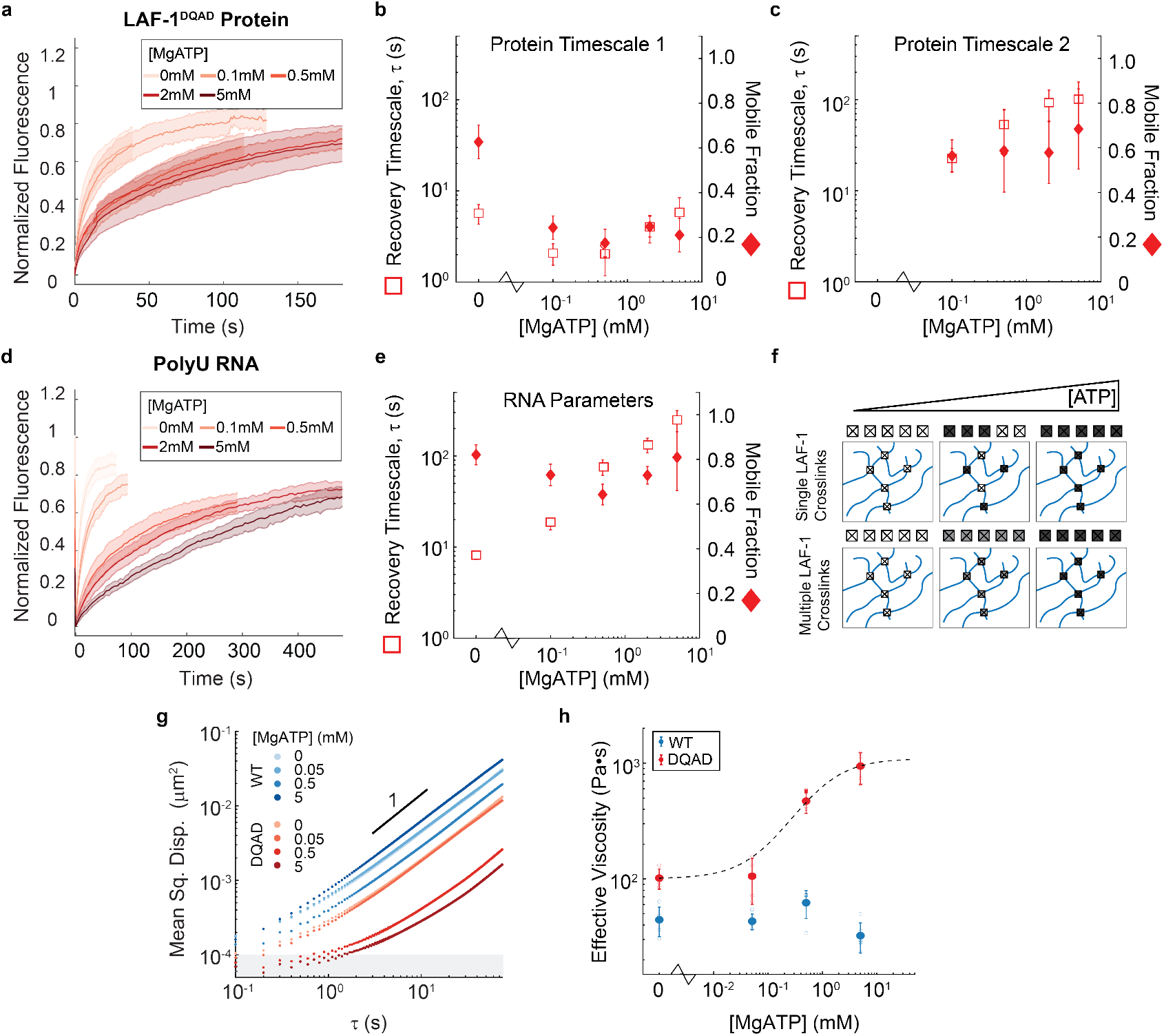
Titratable Crosslink Strength Through ATP Concentration. a) FRAP experiments for labelled LAF-1^*DQAD*^ in condensates containing PolyU RNA and varying concentrations of MgATP show a sharp transition to slowed dynamics above 0.1 mM ATP. b) and c) Recovery timescale (open square) and mobile fraction (closed diamond) fit parameters for the first (b) and second (c) timescales of LAF-1^*DQAD*^ protein FRAP as a function of MgATP concentration. Consistent recovery timescales are observed for the first timescale while the second, longer timescale increases with ATP concentration. d) Titrating ATP results in tunable FRAP recovery dynamics for PolyU RNA in LAF-1^*DQAD*^ condensates. e) Recovery timescale (open square) and mobile fraction (closed diamond) fit parameters of FRAP curves from (d) as a function of MgATP concentration. A monotonic increase in PolyU recovery timescale (*τ*) is observed with increasing ATP concentration. f) Titrating ATP concentration with the LAF-1^*DQAD*^ variant results in tunable network properties via tunable average crosslink strengths. If a single protein can form an RNA crosslink, titrating ATP will increase the overall number of long-lived crosslinks (Top Row). If the crosslinks are formed by multiple LAF-1^*DQAD*^ proteins, then titrating ATP concentration will result in a distribution of crosslink lifetimes with an increasing average lifetime (Bottom Row). g) MSD as a function of time-lag for 100 nm fluorescent tracer particles in LAF-1^*WT*^ and LAF-1^*DQAD*^ condensates containing PolyU RNA of average length 310 nts and varying concentrations of MgATP. LAF-1^*WT*^ and LAF-1^*DQAD*^ experience opposite responses to increased ATP concentration. h) Effective viscosity for condensates containing LAF-1^*WT*^ or LAF-1^*DQAD*^, PolyU RNA of an average length of 310nts, and the indicated concentration of MgATP. Error bars are the standard deviation over individual droplets whose effective viscosities are shown in open circles. The dotted line indicates a hyperbolic fit to the data, reminiscent of LAF-1^*WT*^ ‘s ATPase Michaelis-Menten fit.

The increase in the second timescale of LAF-1^*DQAD*^ FRAP was expected to correspond to an increase in RNA FRAP recovery timescale. We also conducted FRAP experiments as a function of ATP concentration with labelled full length PolyU in LAF-1^*DQAD*^ condensates (Figure 4b). These curves were fit to a single exponential, and we extracted the fit timescale and mobile fraction depicted as a function of ATP concentration (Figure 4d). The mobile fraction is relatively insensitive to ATP concentration, however the RNA recovery timescale monotonically increases as function of ATP concentration. This RNA recovery timescale-ATP concentration relationship strongly resembles the relationship and magnitude of the LAF-1^*DQAD*^ protein’s longer recovery timescale. This is consistent with a model whereby increasing ATP concentration results in an increase in the fraction of LAF-1^*DQAD*^ engaged in a clamp like state along the ssRNA. An increase in clamped proteins results in enhanced crosslinking within the system, which then results in slower average protein-RNA complex mobility (Figure 4f).

If ATP-induced crosslinking was increasing effective macromolecular sizes, we anticipated that such ATP-dependent changes in biomolecular mobility would translate to material properties of the droplets. We performed single particle tracking of beads embedded in the LAF-1 condensates, using an intermediate RNA length of roughly 310 nucleotides and a range of ATP concentrations. LAF-1^*WT*^ condensates experience a slight increase in MSD as ATP increases, likely primarily due to ATP’s hydrotropic properties as opposed to any active transport (Figure 4g, Figure S4) [33]. In contrast, beads within LAF-1^*DQAD*^ condensates experience a decrease in dynamics as ATP concentration increases, with a sharp drop in MSD at ATP concentrations above 50 *µ*M Figure(4g). As all MSD curves were roughly linear in the long time limit, we extracted effective viscosities for LAF-1^*WT*^ and LAF-1^*DQAD*^ condensates (Figure 4h). Upon increasing ATP concentration, we observe a progressive drop in effective viscosity for LAF-1^*WT*^ condensates while LAF-1^*DQAD*^ droplets exhibit a transition from low effective viscosity to high effective viscosity, reminiscent of the behavior seen in the FRAP data (Figure 4h. No heterogeneity is seen in the tracking data indicating either crosslink strength is homogeneously increasing or crosslink spacing is well below the bead size. The behavior in the LAF-1^*DQAD*^ effective viscosity data resembles a hyperbolic relationship between ATP and effective viscosity, as would be expected if the ATP response followed the ATP binding thermodynamics of LAF-1. Upon fitting the LAF-1^*DQAD*^ effective viscosity data to a Hill equation with Hill coefficient of 1, the effective dissociation constant is 0.89mM (+/-0.83mM), consistent with previously reported DEAD-box ATPase K_*M*_ data for other DEAD-box helicases in the presence of RNA as well as the K_*M*_ for the LAF-1^*WT*^ ATPase reaction (Figure S1) [34]. Taken together, this suggests that ATP binding by the LAF-1^*DQAD*^ protein can be utilized to construct condensates with desired RNA mobilities and material properties. Additionally, these results demonstrate that appreciable changes in condensate properties can be achieved with only a small proportion of proteins in a clamp-like state. This is the case in biological systems where any particular mutant DEAD-box helicase composes a minority of the total proteins in a given condensate, yet substantial effects on condensates are still observed with DQAD mutant helicases [11, 16, 35].

## 3 Conclusion

DEAD-box helicases have emerged as critical regulators of biomolecular condensates [6]. Here we contribute to a mechanistic understanding of how modulating DEAD-box helicase-RNA complex lifetimes changes condensate microstructure and dynamics. We show that the DEAD-box helicase DQAD amino acid substitution enables protein-mediated crosslinking of RNA in the presence of ATP. This has consequences for both protein and RNA diffusion within the condensed phase as well as the condensate material state. There are also strong indications that the system approaches a gel transition as RNA length is increased to lengths comparable to eukaryotic mRNA. Lastly, we demonstrate this crosslinking effect can be harnessed to produce condensates with desired properties in accordance with nucleotide concentration. The range over which the system exhibits sensitivity to ATP concentration is highly relevant for biological systems and suggests the LAF-1^*DQAD*^ system can be harnessed for engineering biomaterials in vivo and in vitro.

Our work shows that amino acid substitutions which enhance helicase-RNA interactions can crosslink condensate RNA. In addition to helicase mutation, adaptor proteins may elicit helicase clamp-like states to produce crosslinking in cellular condensates [36–38]. DEAD-box helicases that have longer lived protein-RNA interactions as part of their wildtype activity may also exhibit crosslinking behavior as part of their function, and be fluidized by adaptors which accelerate progress through their ATPase cycle [6, 7, 10]. It may also be possible to elicit a clamp like state via pharmacological methods to crosslink a target biomolecular condensate containing DEAD-box helicases [39, 40]. Importantly, our data suggest a general mechanism for regulating and tuning biomolecular condensate dynamics via nonequilibrium molecular interaction lifetimes, extending beyond DEAD-box helicases.

In addition to coupling ATP binding and hydrolysis to RNA binding and release, DEAD-box helicases utilize this ATPase cycle to unwind duplexed RNA and remodel protein-RNA complexes [10]. This unwinding activity should have implications for condensate dynamics as high degrees of base-pairing decrease biomolecular mobility within condensates [5]. Here we show RNA binding has significant effects, but the interplay between crosslinking and unwinding and their integrated effects on condensate dynamics is unclear. Thus our results demonstrate the need to decouple RNA binding and unwinding activity when considering the effects of DEAD-box helicase mutations and ATP analogs on observed condensate phenotypes. Future work will focus on the involvement of dsRNA and unwinding activity in modulating condensate dynamics to produce a complete picture for DEAD-box helicases and their role as master regulators of biomolecular condensates.

## 4 Methods and Materials

### 4.1 LAF-1 Cloning, Expression, and Purification

The LAF-1 protein coding sequence was ordered as a gBlock Gene Fragment (TM) and inserted into a pET28a(+) vector from Novagen. The LAF-1 mutants E398A (DAAD) and E398Q (DQAD) were developed using around the horn site-directed mutagenesis of the wildtype sequence and similarly inserted into pET28a(+) vectors. Each construct contained a C-terminal His_6_-tag.

Each of the LAF-1 variants was transformed, expressed, and purified as described previously [19]. Briefly, each variant recombinantly expressed and purified from BL21(DE3) cells. The recombinant proteins were then purified using nickel affinity chromatography and heparin affinity chromatography. Purified protein was then dialyzed into storage buffer (20 mM Tris pH 7.4, 1M NaCl, 10% v/v glycerol, 2 mM DTT), flash frozen in liquid nitrogen, and stored at -80°C until use. See supplementary text for more information.

Before use, protein aliquots were thawed at room temperature and buffer exchanged into high salt buffer (20 mM Tris pH 7.4, 1M NaCl, 1 mM DTT) using a 10 kDa cutoff centrifugal filter.

### 4.2 RNA Fragmentation and Capillary Electrophoresis

The ssRNA analog Polyuridylic acid (PolyU) was aquired from Sigma Aldrich and stored as a stock at 10 mg/mL in nuclease free water. Heat and magnesium was used to generate shorter RNA fragments from this PolyU sample, using the NEBNext Magnesium Fragmentation Module. Following fragmentation and reaction quenching, the RNA was isolated and free nucleotides removed using the Monarch RNA Cleanup Kit. RNA was eluted using 14.5 *µ*L of nuclease free water, resulting in RNA concentrations of between 300 and 750 ng/*µ*L. To verify fragment size distributions, capillary electrophoresis was performed on an AATI Femto Pulse.

### 4.3 Fluorescent Labelling

LAF-1 protein was fluorescently labelled using Dylight-633 NHS Ester during purification, following elution from the heparin column. The protein was dialyzed into labelling buffer (100 mM phosphate buffer pH 7.4, 500 mM NaCl, 2mM DTT, 1% v/v glycerol), concentrated to roughly 1 mg/mL, and labelled according to the manufacturers specifications. Excess dye was removed using Pierce dye removal columns. The protein was then dialyzed into storage buffer and then flash frozen. During use, labelled protein composed roughly 1% of the overall protein sample.

PolyU RNA was 3’-end labelled with fluorescein as described in Zearfoss et al. (2012) [41]. For labelling fragmented RNA, smaller elution volumes were used during the RNA cleanup step in order to provide the recommended starting concentrations for 3’-end labelling.

### 4.4 Fluorescence Microscopy and Fluorescence Recovery After Photobleaching

Fluorescence imaging was performed on an Nikon Ti-E inverted microscope equipped with an Andor iXon 897E EM-CCD camera, a Yokugawa CSU-X1 Spinning Disk Confocal Scan Head, and an Andor FRAPPA photomanipulation system. A 100x Nikon objective with a numerical aperature of 1.4 was used for all fluorescence imaging. LAF-1 condensates were formed in buffer containing 20 mM Tris pH 7.4 and 200 mM NaCl, along with 0.075 mg/mL PolyU RNA of various lengths and variable nucleotide concentration. Droplets were incubated for 8 minutes at room temperature and then transferred into a PDMS chamber adhered to a slide and pre-passivated with 2% w/v Pluoronic F-127. For FRAP experiments, a spot size of 3 pixels, corresponding to roughly 0.42 *µ*m, was used for photobleaching.

Analysis of FRAP timecourses was performed using custom packages written in MATLAB and incorporated feature finding scripts from Maria Kilfoil’s MATLAB Point Tracking Software (Supplementary Information).

### 4.5 Point Tracking Microrheology

100nm Diameter carboxy-coated fluorescent microspheres were obtained from ThermoFisher and passivated using mPEG-750 as described in Valentine et al. (2004) [42]. LAF-1 condensates were formed in buffer containing 20 mM Tris pH 7.4 and 200 mM NaCl, along with 0.075 mg/mL PolyU RNA of various lengths, variable nucleotide concentration, and the passivated fluospheres. Droplets were incubated for 8 minutes at room temperature before being briefly centrifuged and then transferred into a PDMS chamber adhered to a slide and pre-passivated with 2% w/v Pluoronic F-127. Condensates larger than 5 *µ*m and containing more than five beads were imaged using a custom built fluorescence microscope equipped with a 561 nm Coherent laser, 100x Nikon 1.45NA oil objective, and an Andor iXon Ultra EMCCD detector. Image series were taken at a framerate of 100 milliseconds using *<* 20 mW laser power.

Feature identification, filtering, and trajectory construction was performed using a MATLAB software package from Maria Kilfoil’s research group [43, 44]. Trajectory analyses such as mean squared displacement, Van Hove correlation, non-Gaussian parameter, and velocity autocorrelation were performed using custom scripts in MATLAB (Supplementary Information).

## 5 Acknowledgements

We would like to thank Peter Foster for helpful discussions throughout this project, Yoon Jung for assembly and maintenance of our in-house fluorescence microscope, and James White for reviewing the manuscript. We would also like to thank Wendy Salmon, Brandyn Braswell, and the W.M. Keck Microscopy Facility for access, training, and expertise in conducting the confocal fluorescence microscopy experiments. We also thank the MIT Biology BioMicro Center for assisting with the capillary electrophoresis experiments. This work was supported in part by the Koch Institute Support (core) Grant P30-CA14051 from the National Cancer Institute, a graduate fellowship from MathWorks Inc., Sloan Foundation Grant G-2021-16758, and the Moore Foundation.

## Supplementary Information

## 1 Extended Methods

### 1.1 Molecular Cloning

The coding sequence of C. elegans LAF-1 with a C-terminal His_6_ tag was synthesized by Genscript and subsequently inserted into a pET28a(+) vector (Novagen) following PCR amplification with primers 1 and 2, and restriction digest with BamHI and NcoI (Table 1).

**Table 1:**
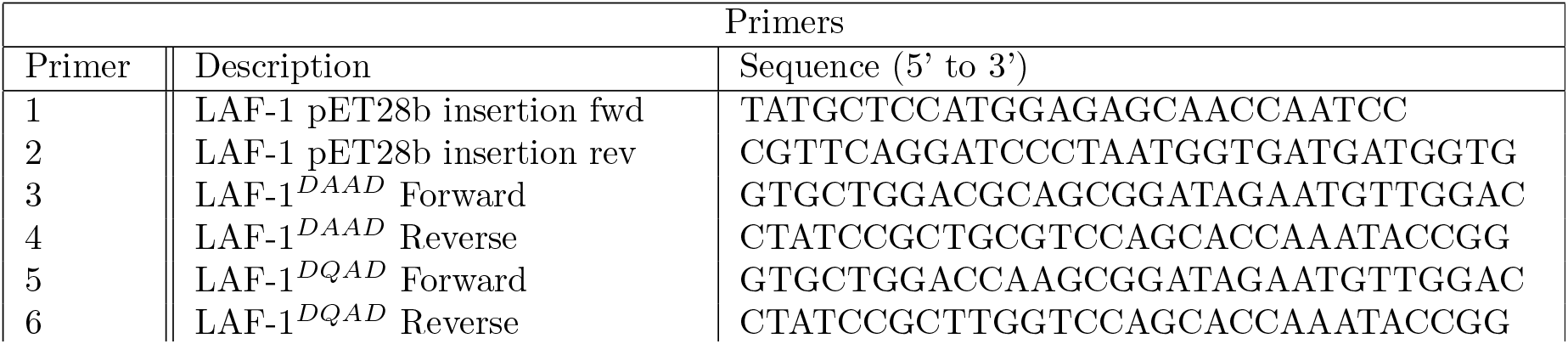
Primer sequences

The mutants E398A (LAF-1^*DAAD*^) and E398Q (LAF-1^*DQAD*^) were developed using around the horn site-directed mutagenesis with primers 3,4 and 5,6, respectively (Table 1). The mutations were confirmed and each mutant construct was subsequently transferred into a fresh pET28a(+) vector.

### 1.2 Recombinant Protein Expression and Purification

LAF-1 was expressed and purified as in Elbaum-Garfinkle et al. (2016). The LAF-1 coding sequence with a C-terminal 6x-Histidine tag was transformed into BL21(DE3) cells on LB Agar plates containing 50ug/mL Kanamycin for selection. Single colonies were transferred to 5mL starter cultures of LB media containing 50ug/mL Kanamycin and grown at 37°C overnight. Starter cultures were transferred to 1L LB Kanamycin media the next morning and grown at 37°C until the OD_600_ was between 0.2 and 0.4. For induction, 1mM IPTG was then added to the cultures and the flasks were transferred to room temperature for an additional 16 hours. Cells were spun down at 8,000 x g, 4°C for 5 minutes, the supernatant removed, and pellets stored at -20°C.

On the day of purification, 1L cell pellets were thawed and resuspended in 50 mL of Lysis Buffer (50 mM Tris pH 7.4, 500 mM NaCl, 10 mM Imidazole, 10% v/v glycerol) containing 14 mM betamercaptoethanol (BME) and an EDTA-free protease inhibitor cocktail tablet (Roche). Lysozyme was added to 100 *µ* g/mL and the mixture was incubated at room temperature for 20 minutes. The lysate was then sonicated on ice with a tip sonicator at 15% amplitude for 5 minutes â€” 15 seconds on 30 seconds off, followed by centrifugation at 20,000 x g for 30 minutes at room temperature. The clarified supernatant was run on a HiTrap Chelating column (GE LifeSciences) charged with 500 mM nickel sulfate and pre-equilibrated with 30 mL of nickel wash buffer (50 mM Tris pH 7.4, 500 mM NaCl, 20 mM Imidazole, 14 mM BME, 10% v/v glycerol). The column was then washed with 50 mL of nickel wash buffer before the sample was eluted with 30 mL of nickel elution buffer (50 mM Tris pH 7.4, 500 mM NaCl, 250 mM imidazole, 14 mM BME, 10% v/v glycerol) in 500 *µ*L fractions. Fractions were tested for their absorbance at 260 and 280 nm and fractions of interest were analyzed on an SDS-PAGE gel before being combined.

The nickel eluate was then diluted 5x with Heparin loading buffer (20 mM Tris, 50 mM NaCl, 2 mM DTT, 1% v/v glycerol) and loaded onto a HiTrap Heparin column (GE LifeSciences) which had been preequilibrated with 30 mL of Heparin loading buffer. Prior to dilution with Heparin loading buffer, the sample was diluted with Nickel Elution buffer to less than 1 *µ*M such that the LAF-1 protein would not undergo phase separation in the Heparin loading buffer. The loaded column was washed with 50 mL of Heparin loading buffer and the sample was then eluted with 30 mL of Heparin Elution Buffer (20 mM Tris, 1 M NaCl, 2 mM DTT, 1% v/v glycerol). Fraction absorbance was read at 260 and 280 nm, fractions of interest were analyzed on an SDS-PAGE, and fractions containing recombinant LAF-1 were pooled together.

For storage, the sample was dialyzed twice in 500 mL of Storage Buffer (20 mM Tris pH 7.4, 1M NaCl, 10% v/v glycerol, 2mM DTT). Samples were concentrated to a protein concentration of between 8 and 12 uM and then 50 *µ*L aliquots were flash frozen in liquid nitrogen. Aliquots were stored at -80°C.

All LAF-1 variants were purified as above. The LAF-1^*DQAD*^ variant copurified strongly with RNA during the nickel affinity purification step, but the A260/A280 ratio dropped to normal levels following heparin affinity purification.

### 1.3 Protein Labelling

Protein samples were labelled using DylightTM 488 NHS-ester or DylightTM 633 NHS-ester (ThermoFisher). Prior to labelling, the protein was dialyzed twice into 500 mL of labelling buffer (100 mM phosphate buffer pH 7.4, 500 mM NaCl, 2mM DTT, 1% v/v glycerol). Protein was then concentrated to roughly 1 mg/mL. Labelling efficiency was higher for Dylight 633 NHS-ester under these conditions. Excess dye was removed using PierceTM Dye Removal Columns. The sample was then dialyzed into Storage Buffer as above and flash frozen in liquid nitrogen before storage at -80°C. Labelled protein was used at 1% of total protein sample during fluorescence experiments.

### 1.4 RNA Labelling

3’-end labelling of PolyU RNA with fluorescein was achieved according to the protocol of Zearfoss et al. (2012) [1].

10 *µ*g of PolyU RNA was oxidized in a 50 *µ*l solution containing 100 mM Sodium Acetate pH 5.2 and 100 *µ*M sodium periodate (NaIO_4_). The reaction was allowed to proceed for 90 minutes at room temperature. The RNA was then precipitated with 2.5 *µ*l of 5 M sodium chloride (NaCl), 1 *µ*l of 20 mg/mL glycogen, and 100 *µ*l of 200 proof ethanol. The mixture was incubated at -20°C for 20 minutes and then centrifuged at 16,000 x g for 25 minutes at 4°C. The supernatant was removed and 50 *µ*l of a fluorescein-5-thiosemicarbazide (FTSC) labelling solution (100 mM sodium acetate pH 5.2, 1.5 mM FTSC) was added. The labelling reaction proceeded overnight at 4°C.

### 1.5 Imaging Chamber Preparation

Polydimethylsiloxane (PDMS) chambers were prepared by mixing 50g of Sylgard 184 Elastomer Base (Dow Chemicals, DC4019862) with 5g of Curing Agent. The mixture was poured into a 150mm x 10mm untreated cell culture dish and degassed under vacuum. The degassed mixture was then cured at 90°C for 90 minutes.

The day before imaging, a 3 mm Integra Miltex Biopsy Punch) was used to create wells in the PDMS surface. The surface of the PDMS was cleaned with Scotch Magic tape, and the PDMS was bonded to the surface of a plasma-cleaned 24×55mm cover slip by incubation on a hotplate for 45 minutes. 2% w/v Pluronic F127 (Millipore Sigma) solution was used to passivate the wells by overnight incubation at 16°C in a humidified chamber.

Before use, the Pluronic F127 solution was removed and each well was washed 4 times with assay buffer.

### 1.6 Bead Passivation

Red-fluorescent carboxy-coated FluoSpheres^*T M*^ with a 0.1 um diameter were obtained from ThermoFisher (F8801) and passivated with amine-terminated methoxy-poly(ethylene glycol) (mPEG-NH_2_) containing an average PEG size of 750 kDa, according to the protocol of Valentine, M.T. et al. (2004).

The 100 nm carboxy-coated microspheres were diluted to 4 *∗* 10^12^ particles per mL and tip sonicated (10% amplitude, 5 seconds on, 5 seconds off, 30 seconds total time). 2mL of the diluted bead solution was dialyzed into 300 mL of 100 mM 2-(N-norpholino)ethanesulfonic acid (MES) at pH 6.0 for 2.5 hours. The dialysis bag was then rinsed with deionized water before being transferred to a 500 mL solution containing 100 mM MES, 15 mM 1-[3-(dimethylamino)rpropyl]-3-ethylcarbodiimide (EDC), 5 mM N-hydroxysuccinimide (NHS), and a 10x excess of mPEG-NH_2_. The reaction was allowed to proceed for 45 minutes. The dialysis bag was then transferred to 500 mL of borate buffer (50 mM boric acid, 36 mM sodium tetraborate) at pH 8.5 containing 5 mM NHS, and a 10x excess of mPEG-NH_2_ and the dialysis continued for at least 8 hours. This dialysis was repeated 2 more times, and then the dialysis cassette was transferred to pure borate buffer and dialyzed for 4 hours. Finally, the bead solution was removed from the dialysis cassette and stored at 4°C.

### 1.7 Bead Tracking

LAF-1 droplets were formed in the presence of passivated 100nm fluospheres, with microspheres at a concentration of roughly 1/2000 of the stock concentration. The sample was allowed to equilibrate for 5-10 minutes before being transferred to a PDMS chamber passivated with 2% w/v PF-127 (see above). Droplets larger than 5 *µ*m and containing greater than or equal to 5 beads were imaged with a frame-to-frame interval of 100 msec using a 561 nm laser. Experiments were conducted at 21°C +/-1°C.

Post-processing of images was conducted in ImageJ while feature finding and trajectory construction were performed using Maria Kilfoil’s MATLAB implementation of the Crocker-Grier algorithm [2, 3]. Analysis of trajectories was performed using custom MATLAB scripts discussed below.

### 1.8 Fluorescence Recovery After Photobleaching

For FRAP analysis, bleach spot centroids were identified by using feature finding scripts from Maria Kilfoil’s microrheology software applied to thresholded and gradient processed images performed in MATLAB. Normalization was performed as previously described [4]. Average fluorescent intensity within the bleached region at time t (*(C*(*r, t*))) was normalized relative to the average fluorescent intensity within the bleached region immediately following bleaching (*(C*(*r*, 0))) and the average fluorescence intensity far from the bleached region at time t (*(C*_*ref*_ (*t*))) to produce the normalized fluorescence intensity (*C*^*∗*^(*t*)):

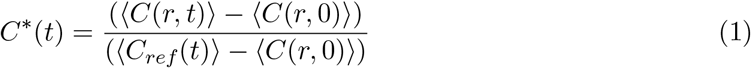

Error bars presented in normalized fluorescence intensity traces are the standard deviation calculated over all droplet FRAP traces measured for that condition.

Normalized fluorescence intensity time traces were then fit to single exponentials:

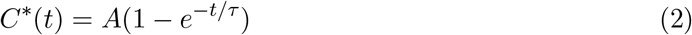

where the A parameter defines the mobile fraction and the *τ* parameter defines the recovery timescale and is related to macromolecular diffusion.

For the LAF-1^*DQAD*^ protein FRAP when ATP is present, recovery dynamics are better described by a double exponential, suggestive of two protein diffusion modes within the condensed phase:

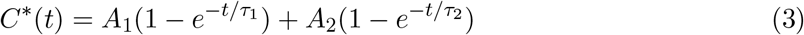

where *A*_1_ and *τ*_1_ are the mobile fraction and recovery timescale, respectively, of the faster diffusion mode and the *A*_2_ and *τ*_2_ are the mobile fraction and recovery timescale of the slower diffusion mode, respectively. At low or zero ATP concentrations, we cannot justify two timescales in the protein FRAP fits for LAF-1^*DQAD*^. Similarly, a single exponential is sufficient to describe the dynamics of the wildtype protein, suggesting that protein bound to RNA releases on a timescale faster than the protein or RNA diffusion timescale.

Fits were performed to individual droplet FRAP traces as well as to averaged FRAP traces across all droplets for that conditions. The mean and standard deviation of the individual droplet fit parameters is displayed in Supplementary Table 3. The fit parameters for the fit to the mean FRAP traces are also displayed in Supplementary Table 3. Error bars in FRAP parameter plots display the standard deviation of parameters for fits to individual droplets.

### 1.9 ATPase Assays

ATPase assays were conducted using the Abcam ATPase/GTPase assay kit (ab272520). Assays were conducted using a LAF-1 protein concentration of 500nM in 20mM Tris pH 7.4, 200mM NaCl, and 1mM DTT. Reactions were monitored at several time-points over 90 minutes using the indicated concentration of MgATP. At each time-point, 40 *µ*l of the reaction was removed and mixed with 200 *µ*l of malachite green assay reagent. The sample was then transferred to a 96-well plate and the absorbance of the sample at 620nm was read out on a Tecan Infinite M200 plate reader. Each condition was tested in three independent assays. Absorbance measurements were correlated to phosphate release and ATP turnover using standards provided in the kit. Rates were then extracted by linear fits of the free phosphate production over time.

ATPase rates as a function of ATP concentration were fit to a Michalis-Menten equation:

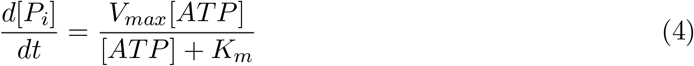

Fitting was performed in MATLAB, using the nlinfit function from the Statistics and Machine

Learning toolbox. Fit parameters extracted along with their 95% confidence intervals:

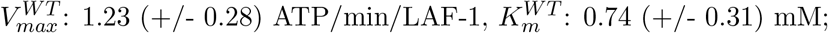

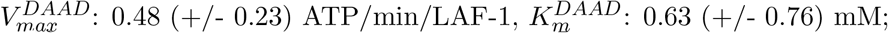

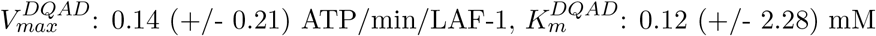

### 1.10 Partition Coefficient

Fluorescence micrographs of condensates containing fluorescently labelled LAF-1 and labelled PolyU RNA were thresholded and segmented using elements of the Image Processing Toolbox in MATLAB. Condensate boundaries were defined relative to the protein fluorescence channel, and subsequently the distribution of pixel fluorescence intensities for both protein and RNA channels inside a particular condensate was measured to extract a mean pixel intensity. The mean pixel intensity outside all condensate boundaries within a field of view away from the coverslip was then measured to estimate the dilute phase fluorescence intensity. Droplets below a particular size were removed from consideration. Partition coefficient was then calculated for each droplet by taking its mean fluorescence intensity and dividing by the dilute phase fluorescence intensity.

### 1.11 Trajectory Analysis

#### 1.11.1 Mean Squared Displacement

Time-ensemble-averaged mean Squared Displacement (TMSD) of a condition was calculated by averaging all mean squared displacements over time window *τ* (*r*(*t* + *τ*) *− r*(*t*)) across all windows of size *τ* for a single bead (n) and all beads (m):

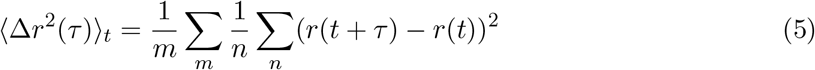

TMSDs for most of the conditions tested were diffusive in the long time limit ((Δ*x*^2^(*τ*)*) ∼ τ*), consistent with fluid-like behavior, and the subdiffusive behavior at short times was well described by accounting for a constant noisefloor additive term in fitting the bead MSDs. This noisefloor was measured by calculating the TMSDs of beads adhered to a coverslip, whose TMSD should then indicate noise contributions from tracking and stage vibration. The measured noisefloor was 1e-4 *µm*^2^.

#### 1.11.2 Effective Viscosity and Stokes-Einstein Relation

For those droplets with bead MSDs that exhibited diffusive scaling with time-lag, the droplet TMSD was fit to a linear fit in MATLAB:

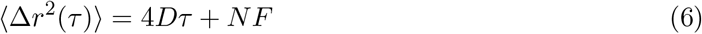

Where *τ* is the time-lag in seconds, D is the diffusion coefficient, and NF is the noisefloor. Effective viscosities (*η*_*eff*_) were then calculated using the Generalized Stokes Einstein Relation (GSER):

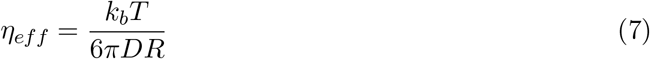

R is the radius of the particle (50 nm), *k*_*b*_ is the Boltzmann constant, and T is the temperature in Kelvin. Experiments were conducted at 21°C +/-1°C (294K).

For Figure 4h, the effective viscosity as a function of ATP concentration data for the LAF-1^*DQAD*^ variant was fit to a rectangular hyperbola with a Hill coefficient of 1 and a scaling term describing the maximal effective viscosity of the system (similar to the form of the Michaelis-Menten equation):

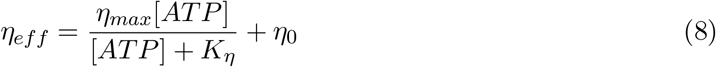

*η*_*max*_ and *K*_*η*_ are analogous to *V*_*max*_ and *K*_*m*_ within the Michaelis-Menten equation, and *K*_*η*_ is the ATP concentration at which half the total transition in effective viscosity is achieved for the RNA length of 310nts. Here an intercept (*η*_0_) is included as the effective viscosity at a concentration of ATP is not expected to be zero. We have not included it as a free parameter in our fitting, and have fixed it at the system’s effective viscosity with 0mM ATP present, 101.7 Pa·s.

Fitting was performed using MATLAB, via the nlinfit function within the Statistics and Machine Learning Toolbox. 95% confidence intervals of each parameter were also extracted using the nlparci function. Starting fit parameters of 500 Pa·s and 0.7mM were used for *η*_*max*_ and *K*_*η*_, respectively. Final fit to the equation above yielded an *η*_*max*_ of 995 Pa·s (95% confidence interval +/-268 Pa·s) and a *K*_*η*_ of 0.89mM (95% confidence interval +/-0.83mM).

#### 1.11.3 Van Hove Histograms

The step size histograms for a particular droplet were calculated across all bead trajectories and subtrajectories of length *τ* :

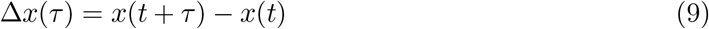

Bead diffusion within a simple fluid exhibit Brownian motion and are expected to have a Gaussian distribution of step sizes.

#### 1.11.4 Non-Gaussian Parameter

To quantify deviation from Gaussian behavior, we can calculate the Non-Gaussian Parameter (NGP) for a particular droplet by considering all trajectories from that droplet and then calculating the ratio of the fourth moment of the displacement distribution to the second moment [5]:

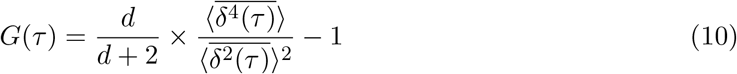

where d is the dimension, 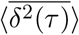 is the TMSD, and the fourth time averaged moment is:

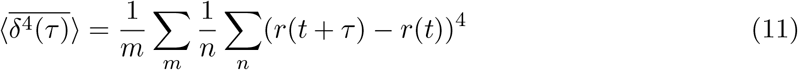

The summations are over m particles and n sliding windows.

Positive values of *G*(*τ*) indicate a deviation from a Gaussian distribution, whereas the NGP approaches zero for Gaussian particle displacement distributions at that particular time-lag.

#### 1.11.5 Velocity Autocorrelation Function

The velocity autocorrelation function (VACF, 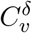) for a droplet can be calculated by considering how velocities of individual beads 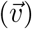 are self-correlated as a function of time separation between velocity measurements (*τ*) [6]:

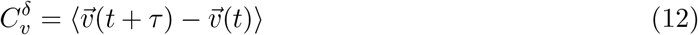

Here velocities can also be defined over a variable time window (*δ*) and are vector quantities.

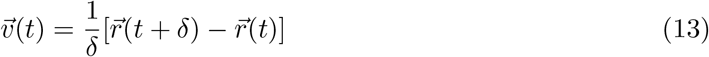

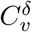 is typically normalized relative to the autocorrelation at *τ* = 0 to allow VACFs using different velocity window sizes (*δ*) to be compared more easily.

The expectation for a Brownian trajectory (as in simple fluids) is for the normalized VACF to start at one and decrease to zero as the window over which the velocities are calculated from overlapping sequences decreases, up until *τ* = *δ*. At this point, the velocities should become uncorrelated and the normalized VACF should remain around zero for *τ > δ*. Viscoelastic materials will typically produce negative VACF peaks around *τ* = *δ*, which then tend to zero at longer *τ*. Active transport of particles will typically manifest as positive VACF values at *τ* = *δ*, which then decay to zero at longer *τ*.

## 2 Supplementary Tables

Table 2: FRAP Fit Parameters for LAF-1 Variants in the Presence of different RNAs, nucleotides, and nucleotide concentrations

Table 3: Averaged Point Tracking Microrheology for LAF-1 Variants in the Presence of different RNAs, nucleotides, and nucleotide concentrations

## 3 Supplementary Figures

**Figure S1:**
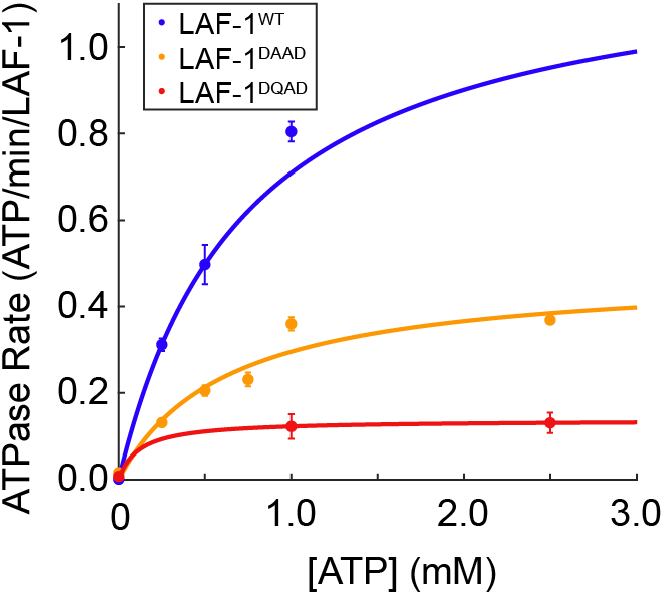
ATPase Activity of the LAF-1 Variants. Mutations within the catalytic core, DAAD and DQAD, exhibit decreased ATPase activity relative to LAF-1^*WT*^. Error bars are the standard deviation across three replicates for each condition tested. Solid lines indicate fits to the Michaelis-Menten equation: 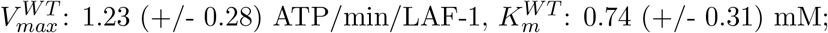 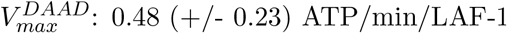 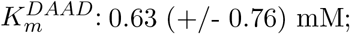 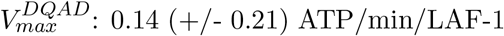 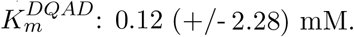

**Figure S2:**
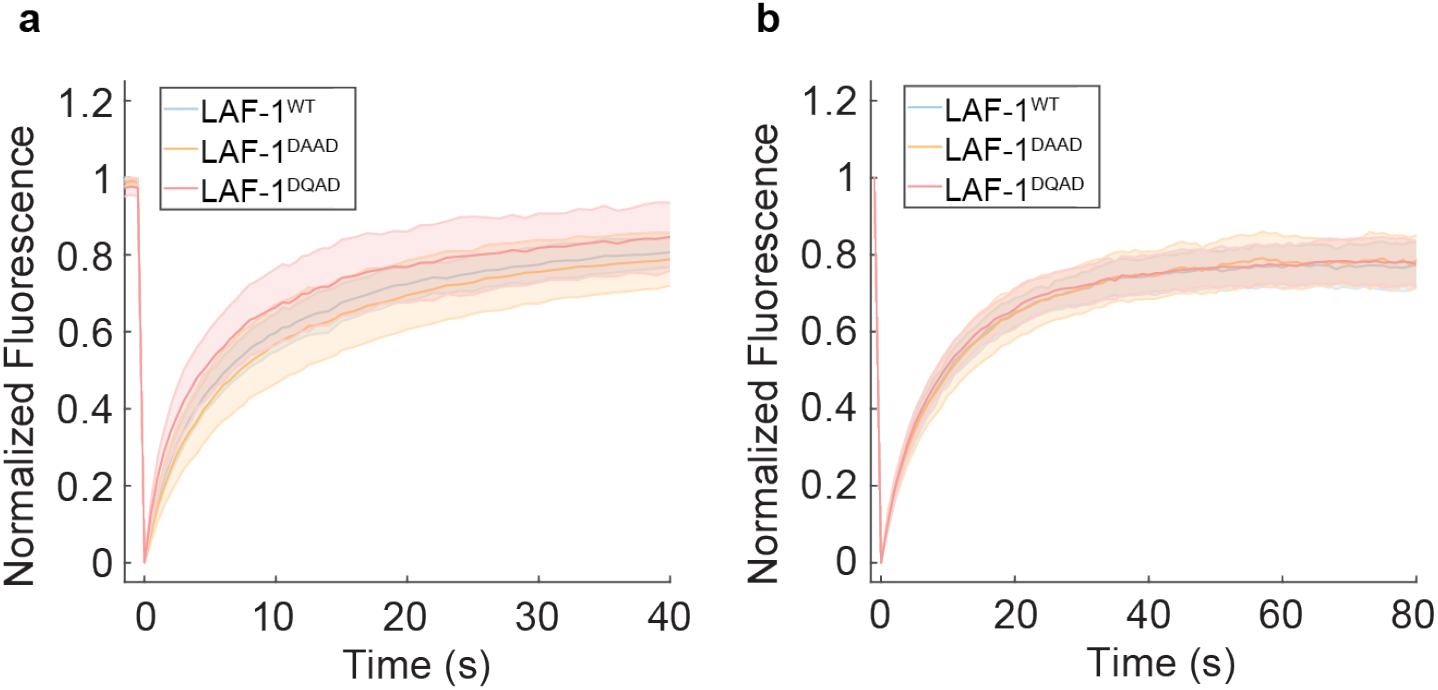
FRAP response between variant condensates is identical in the presence of MgADP. a) FRAP curves for labelled LAF-1 protein variants in LAF-1-RNA-MgADP condensates. b) FRAP curves for labelled PolyU RNA in LAF-1-RNA-MgADP condensates of the indicated LAF-1 variant. Error bars are over all droplets measured for the respective condition.

**Figure S3:**
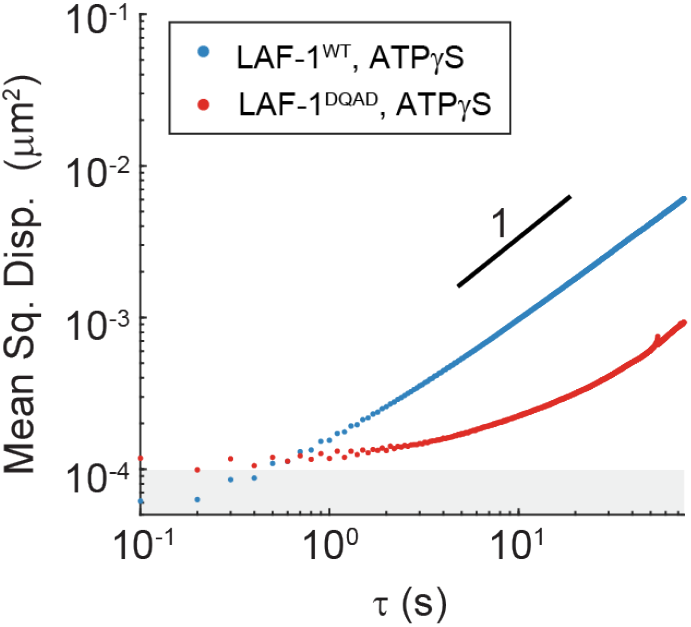
LAF-1^*WT*^ and LAF-1^*DQAD*^ exhibit similar bead tracking dynamics in the presence of ATP*γ*S and PolyU ssRNA to their counterpart systems with ATP and PolyU ssRNA. Noisefloor of 1e-4 *µm*^2^ is shown by the grey rectangle. MSD scaling exponent of 1 is shown by the black bold line.

**Figure S4:**
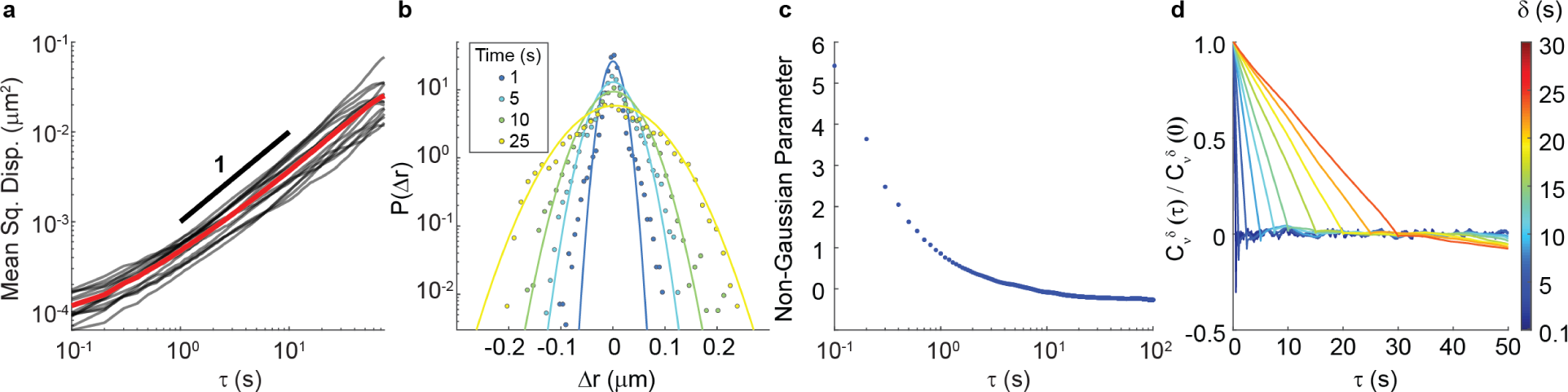
Brownian behavior of beads in LAF-1^*WT*^ condensates containing unfragmented PolyU RNA and 5mM MgATP. a) Individual MSDs (black) and time-ensemble averaged MSDs (red) as function of time-lag (*τ*) for beads in a single LAF-1 condensate containing PolyU RNA and 5mM ATP. MSD of slope 1 is shown by the black solid line. IMSDs and the TMSD show agreement with this diffusive expectation. b) Van Hove histograms for step sizes for the beads in (a) for different times. Step size distributions have good agreement with the Gaussian expectation for Brownian motion at times wherein the bead motion has escaped the noisefloor of approximately 1e-4 *µm*^2^. c) Non-Gaussian Parameter (NGP) (see SI Text) as a function of time-lag for the system. The NGP approaches the Brownian expectation of 0 as the system escapes the noisefloor. No peaks are seen at intermediate timescales. d) Velocity autocorrelation function (VACF) of bead trajectories within the system for different velocity time windows (*δ*) as a function of time-lag (*τ*) between velocity calculations (See SI Text). VACFs decay to 0 as *τ* approaches *δ* and then remain at zero for longer time-lags, consistent with diffusive, Brownian trajectories. Negative peaks at *τ* = *δ* for small *δ* are consistent with noisefloor contributions at short times.

**Figure S5:**
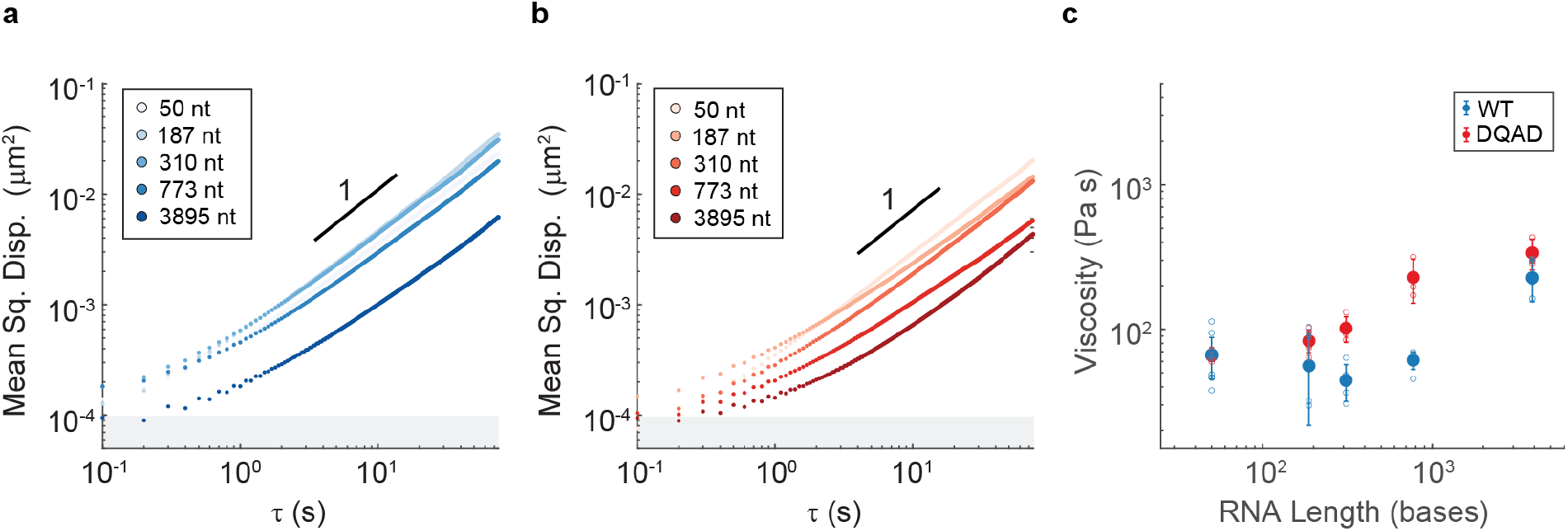
LAF-1^*WT*^ and LAF-1^*DQAD*^ condensates behave similarly as a function of length in the absence of ATP. a) and b) MSD as a function of timelag for 100 nm fluorescent tracer particles in LAF-1^*WT*^ and LAF-1^*DQAD*^ condensates containing PolyU RNA with the indicated average length and 0 mM MgATP. Noisefloor of 10^*−*4^*µm*^2^ is shown by the gray rectangle. MSD scaling exponent of 1 is shown by the solid line. c) Effective viscosity for condensates containing LAF-1^*WT*^ or LAF-1^*DQAD*^ (blue or red, respectively), PolyU RNA of the indicated length, and 0mM ATP, as extracted using the Generalized Stokes-Einstein Relation. Average effective viscosity is shown by the solid circles while individual droplet effective viscosities are shown in open circles. Error bars are the standard deviation of effective viscosity across all droplets measured for that condition. General agreement is seen between the two LAF-1 variants for all RNA lengths tested.

**Figure S6:**
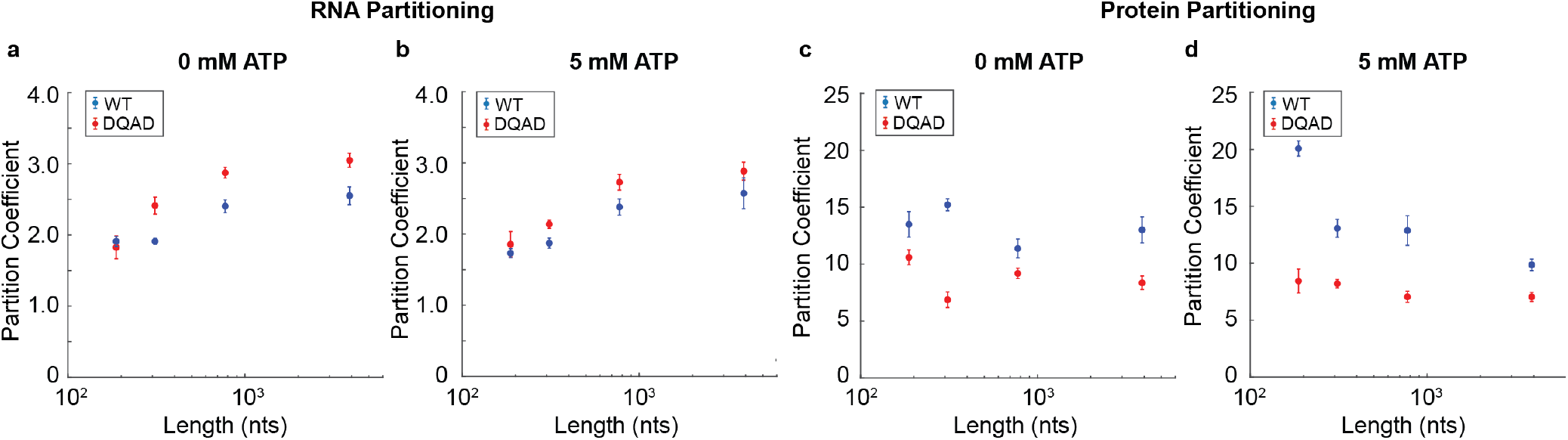
Partition Coefficient as a Function of RNA Length, ATP Concentration, and LAF-1 Variant. a) and b) Partition coefficient of PolyU ssRNA of different lengths labelled with fluorescein in condensates of LAF-1^*WT*^ (blue) and LAF-1^*DQAD*^ (red) with either 0 mM or 5 mM MgATP present in the system, respectively. c) and d) Partition coefficient of LAF-1 protein labelled with Dylight633 in systems containing PolyU of varying lenths for LAF-1^*WT*^ (blue) and LAF-1^*DQAD*^ (red) with either 0 mM or 5 mM MgATP present in the system, respectively. Partition coefficients were calculated using fluorescent signal inside of LAF-1-RNA condensates and fluorescent signal in the dilute phase away from the cover slip. Error bars are the standard deviation across all droplets measured.

**Figure S7:**
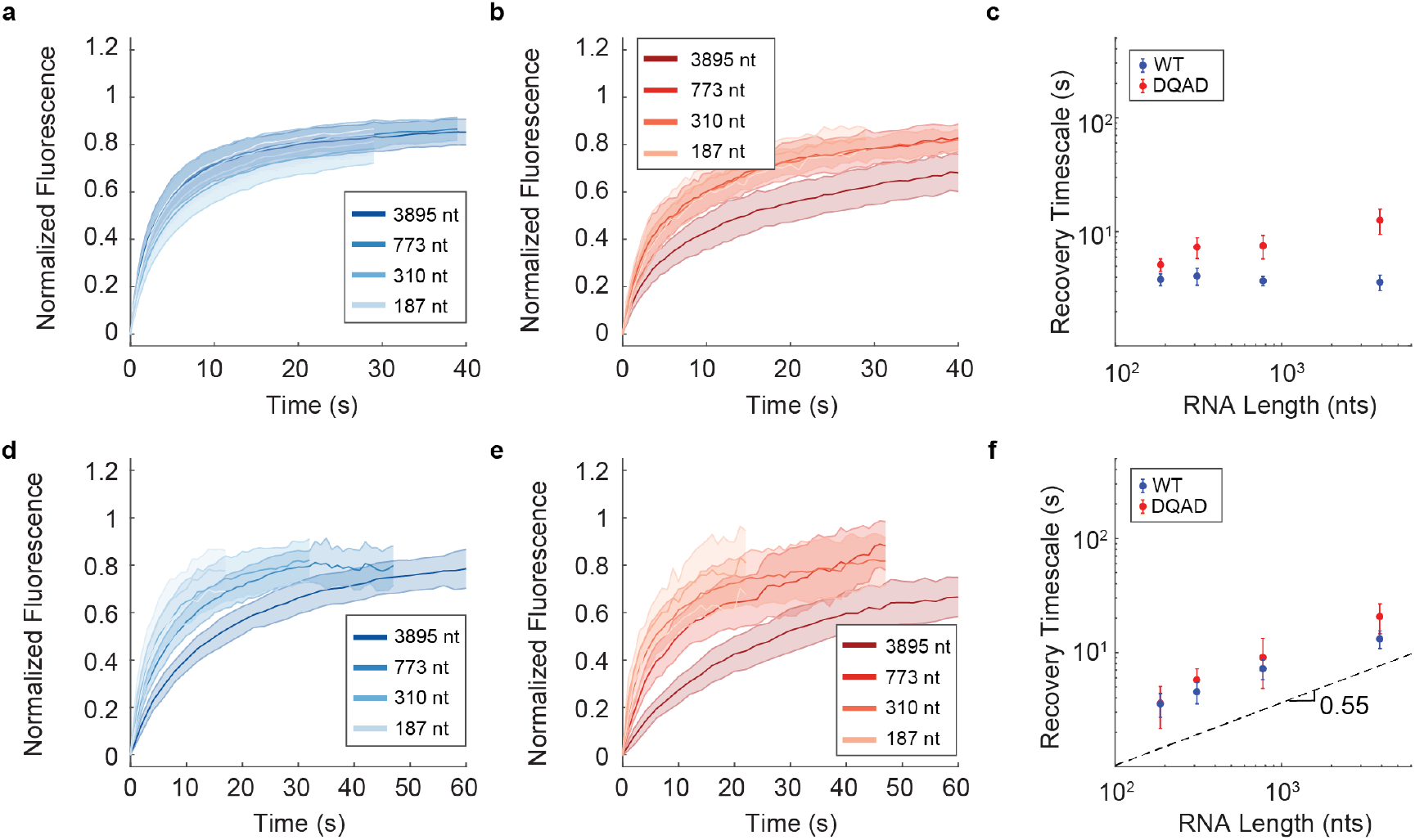
Increased Effective Cluster Size for RNA in LAF-1^*DQAD*^ Condensates. a) and b) FRAP curves for labelled LAF-1 protein in LAF-1^*WT*^ and LAF-1^*DQAD*^ condensates, respectively, with varying PolyU lengths and 5 mM ATP. Error bars are standard deviation in normalized fluorescence intensity measured over all droplets. c) FRAP recovery timescales for LAF-1 protein FRAP from (a) and (b), where both LAF-1^*WT*^ and LAF-1^*DQAD*^ protein recovery was fit to a single exponential. Error bars are the standard deviation in individual droplet fit parameters over all droplets for that condition (Table S2). Good agreement is seen between the LAF-1^*WT*^ and LAF-1^*DQAD*^ recovery timescales, with no significant dependence on RNA length. d) and e) FRAP curves for labelled PolyU RNA in LAF-1^*WT*^ and LAF-1^*DQAD*^ condensates, respectively, with varying PolyU lengths and 5 mM ATP. Error bars are standard deviation in normalized fluorescence intensity measured over all droplets. f) RNA FRAP recovery timescales fit to single exponentials. Error bars are the standard deviation in individual droplet fit parameters measured over all droplets for that condition (Table S2). Good agreement is seen between the two variants and roughly 0.55 scaling is seen between recovery timescale and RNA length, consistent with diffusion coefficient as function of molecular weight and *R*_*g*_.

**Figure S8:**
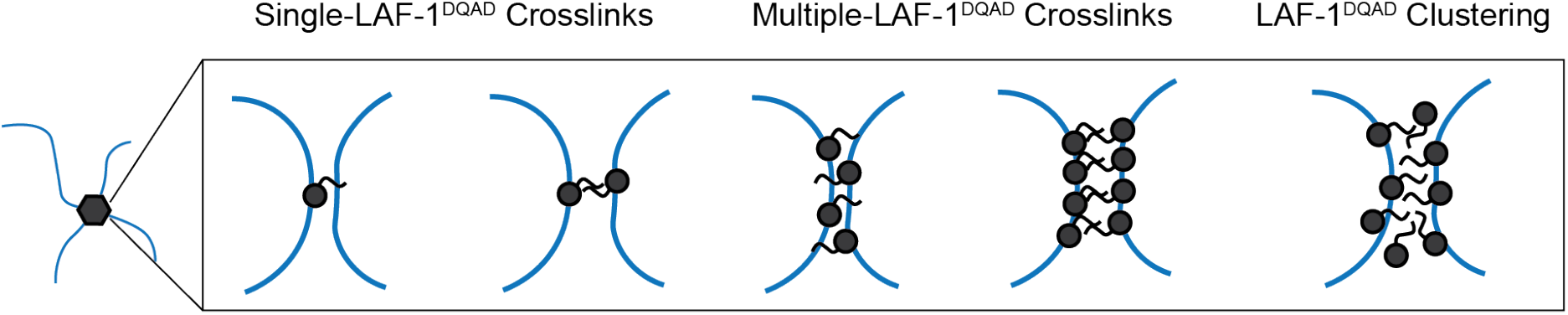
Different Potential Models for LAF-1^*DQAD*^ crosslinking of ssRNA. Black depicts LAF-1 molecules while blue depicts RNA molecules. The black circles represent the LAF-1^*DQAD*^ helicase core while the black squiggle represents the N-terminal disordered domain of LAF-1.

